# Magnetic Affinity-Based Purification of His-Tagged Proteins Using Ni^2+^-Functionalized Nanoparticles

**DOI:** 10.1101/2025.08.04.668490

**Authors:** Carolina Otonelo, Elisa de Sousa, Luciana Juncal, Pedro Mendoza Zelis, Sheila Ons, Claudia Rodríguez Torres, Carla Layana

## Abstract

Recombinant protein purification is essential in many applications across industry, research, and biotechnology. The use of a histidine tag (His6) in affinity chromatography is the gold standard for purification. However, magnetic nanoparticles (MNPs) offer an excellent alternative due to their unique properties, such as rapid separation, superparamagnetism, and a high surface-to-volume ratio that facilitates nanoscale functionalization. MNPs provide a more efficient, faster, cost-effective, and sustainable method compared to traditional chromatography. In this work, we report the development and validation of a magnetic nanoparticle-based platform for the efficient, selective, and scalable purification of His6 recombinant proteins. The synthesized Ni^2+^-functionalized MNPs demonstrated high specificity for histidine-tagged proteins, as shown by the selective retention and imidazole-mediated elution of target proteins both from pre-purified samples and complex *E. coli* lysates. Comparative assays with bovine serum albumin and His6 protein A confirmed the selective affinity of the MNPs for His6. Furthermore, a scaled-up protocol using 4 mL of lysate yielded approximately 0.6 mg/mL of purified protein, doubling the output obtained via conventional immobilized metal affinity chromatography, while preserving purity. The simplicity, efficiency, and cost-effectiveness of this method render it a powerful tool for both analytical and preparative-scale protein purification, particularly in low-resource or high-throughput settings.

## Introduction

In recent years, there has been an increasing demand for large amounts of pure, soluble, and biologically active proteins, driven by their critical role in research, as well as therapeutic, diagnostic, proteomic, biopharmaceutic and industrial applications. However, protein purification from natural sources often falls short in meeting the requirements for quantity, quality, ease of isolation, and economic feasibility. The advent of recombinant DNA technology in the mid-1970s ushered in a new era of biotechnology. Today, with advances in genomics, proteomics, and bioinformatics, the production and purification of recombinant proteins have expanded rapidly (Ana et al., 2022). The ability to selectively recover proteins from complex biological environments is fundamental for the study and use of recombinant proteins. Among the various expression systems available, the bacterium *Escherichia coli* remains the system of choice, thanks to its fast growth, high biomass yield in cost-effective media, well-understood physiology and genetics, and its remarkable ability to produce heterologous proteins at both laboratory and industrial scales (Jia and Jeon, 2016; Morão et al., 2022; Rosano et al., 2019; Selleck and Tan, 2008; Studier, 2014). Although various purification techniques have been developed, such as ultrafiltration, electrophoretic separation, size-exclusion chromatography, ion-exchange chromatography, and others (Du et al., 2022; Hanno et al., 2003; Richard, 2018), in the field of recombinant protein purification, the use of a histidine tag (His6) in affinity chromatography has become the gold standard due to its high specificity and efficiency (Elliott et al., 2020; López-Laguna et al., 2022; Valeria et al., 2020).

Over the past few years, the use of magnetic nanoparticles (MNPs) for protein purification has gained significant attention due to their unique properties, such as rapid separation, efficiency, speed, cost-effectiveness and high surface-to-volume ratio that facilitates their functionalization at the nanoscale (Eivazzadeh-Keihan et al., 2021; Le et al., 2023). In contrast to conventional methods, MNP-based purification enables rapid and selective separation without requiring extensive sample pretreatment, such as centrifugation or filtration, and does not rely on sophisticated equipment. Advancements in surface functionalization techniques have improved the specificity and binding capacity of MNPs, allowing for the efficient capture of target proteins through affinity interactions, such as the interaction of His6 proteins with Ni^2+^-coated MNPs. Additionally, the development of superparamagnetic nanoparticles has enhanced the control and recovery of proteins under external magnetic fields, reducing loss and contamination.

MNPs, typically composed of iron oxide (Fe_3_O_4_), can be functionalized with a variety of ligands, such as metal ions (Ni^2+^, Cu^2+^, Co^2+^, Zn^2+^), antibodies, or polymers (Leonhardt et al., 2023), allowing for selective binding to target proteins or other biomolecules (Liu et al., 2020). The use of MNPs in immobilized metal ion affinity chromatography (IMAC) for the purification of His6 proteins is particularly advantageous due to the strong affinity between histidine residues and metal ions. This results in faster purification times, higher protein yields, and simpler recovery steps compared to traditional column-based chromatography methods. Another advantage of MNPs is their reusability, after purification, the nanoparticles can be washed and regenerated for multiple rounds of protein capture, making them a more cost-effective solution for large-scale operations (Krasitskaya et al., 2022; Ma et al., 2021; Salimi et al., 2017).

Here, we present an efficient protocol for His-tagged protein purification from bacterial cultures using MNP@Ni^2+^ particles synthesized in our lab. We compared this magnetic purification approach with conventional chromatographic methods, and demonstrated that it provides a practical and effective strategy for obtaining proteins suitable for research and industrial applications.

## Materials and Methods

### Synthesis of MNP@Ni^**2+**^

MNPs were obtained using the coprecipitation method. For this, 2.7 g of iron (III) chloride hexahydrate (FeCl_3_.6H_2_O) and 1 g of iron (II) chloride tetrahydrate (FeCl_2_.6H_2_O) were mixed in a 250 ml flask containing 20 ml of water, under magnetic stirring and heated at 60°C. After 5 minutes of incubation, salts were completely dissolved, 5 ml of 25% ammonium hydroxide (NH_4_OH) were slowly added dropwise using a burette, and the solution was incubated for 30 minutes under the same conditions. The resulting magnetite MNPs were washed twice using an 1:1 ethanol:water mixture with the help of a magnet, and finally resuspended in 250 ml of ethanol. Before proceeding with the next steps, the nanoparticles were sonicated for 30 minutes to ensure good dispersion in the solution, and 5 ml were taken for characterization. This washing and sample reservation step for characterization was repeated after each step of the synthesis.

The bare MNPs were coated with silica following a modified Stöber method (Lee et al., 2018), which involves the alkaline hydrolysis of tetraethyl orthosilicate (TEOS) in an ethanol/water mixture in the presence of MNPs. For this, 312 µl of TEOS were added dropwise to the solution and left to incubate for 6 hours at room temperature under stirring. Once MNP@TEOS were obtained, a variation of Minkner’s synthesis (Minkner et al., 2020) was followed, 480 µl of APTES were slowly added to the NP@TEOS suspension and incubated for 36 hours under reflux at 60 °C with stirring. Then, 1 g of isatoic anhydride was added to the solution, maintaining agitation and reflux at 80°C for 12 hours. As the final step, 0.4 g of NiCl_2_ were added, maintaining stirring and reflux at 80 °C for 12 hours.

The resulting MNP@Ni^2+^ were dried using a vacuum pump and stored as a powder at 4 °C until use. The MNP@Ni^2+^ particles exhibit superparamagnetic behavior, with a saturation magnetization (Ms) of 60 emu/g. Considering that the Ms of the bare magnetic nanoparticles (MNPs) is 75 emu/g, this result indicates that approximately 20% of the total mass corresponds to non-magnetic coatings, such as silica and other surface functionalization layers. In addition, dynamic light scattering (DLS) measurements showed that MNP@Ni^2+^ have a positive zeta potential of +(30 ± 2) mV, compared to −(30 ± 2) mV for the bare MNPs, confirming the presence of Ni^2+^ ions on the surface. The hydrodynamic diameter of MNP@Ni^2+^ was found to be (210 ± 10) nm.

## Induced expression of recombinant proteins

The sequence corresponding to domain D of *Staphylococcus aureus* protein A (*GenBank: U54636*.*1*) was synthesized and cloned into the pDEST 14-SpyCatcher vector (4061 bp, *Macrogen*, Jin et al 2021). Cloning at the NcoI site was chosen so that the protein A sequence remains in the reading frame at the amino terminus of Spy Catcher and contains His6. In this way, the expressed protein (His6-SC-ProtA) will have a size of ~21 kDa.

Recombinant protein expression was carried out using *E. coli* BL21 (DE3) cells transformed with the expression plasmid containing the gene of interest. Bacterial cultures were grown in LB medium supplemented with antibiotics at 37 °C with shaking (200 rpm) until reaching an optical density at 600 nm (OD600) of 0.6-0.8.

Protein expression was induced using 1 mM isopropyl β-D-1-thiogalactopyranoside (IPTG), and the cultures were incubated for 3 hours at 37 °C. Bacterial cells were harvested by centrifugation at 5,000 x g for 10 minutes, and the resulting pellet was resuspended in lysis buffer (50 mM Tris-HCl, pH 8.0; 5 mM EDTA, pH 7.4; 1x protease inhibitor cocktail; 0.43 mg/ml lysozyme; 10 mg/ml RNase; 1 U/ml DNase I). The resuspended cells were incubated for 30 minutes with agitation at 4°C, followed by three 10 second pulses of sonication (*Branson Sonifier 250*), interspersed with ice baths to prevent overheating. Cell lysates were clarified by centrifugation at 15,000 rpm for 15 minutes at 4°C. Finally, the supernatants were stored at −20°C until further use.

### Protein purification by column method

The whole purification process was performed on an *AKTA START chromatography system* (*GE, Pittsburgh, USA*). The lysate was loaded onto a 1 ml HisTrap column (*Cytiva*) with a loading volume of 50ml, 7 ml of lysates and 43 ml of binding buffer (20 mM phosphate buffer, 150 mM NaCl, 20mM imidazole). Next, the column was washed with the binding buffer until the absorbance at 280 nm reached a steady baseline. The His6 proteins were eluted with the elution buffer (500 mM imidazole) The eluted proteins were dialyzed with gentle stirring in 1x PBS (pH 7.4) at 4°C for 1 hour twice, with a final overnight step. After imidazole was removed, the purified proteins were further analyzed via SDS-PAGE (see below) and were stored at −20°C until use.

### Protein purification by MNPs

Three different protocols were employed for protein purification using magnetic nanoparticles, depending on the nature of the starting material: (i) purified His6 protein, (ii) clarified *E. coli* lysate, or (iii) large-scale clarified lysate. Unless stated otherwise, all steps were carried out at 4°C.

i. Purified His6 protein For binding assays using a purified His6 protein, 0.6 mg of MNPs were suspended in washing buffer (20 mM Tris-HCl, 0.5 M NaCl, 7.5 mM imidazole, pH 7.5) and incubated with 115□μg of purified His6-SC-ProtA. The final volume and buffer volume were adjusted as needed to allow proper dispersion. The mixture was incubated for 3 hours under gentle agitation to promote binding. After incubation, the sample was placed on a magnetic rack, and the supernatant was removed. The nanoparticles were then washed twice using 150 μL of washing buffer, with each wash followed by magnetic separation. Elution was performed in three steps. The first elution used 250 μL of a weak elution buffer (20 mM Tris-HCl, 0.5 M NaCl, 112.5 mM imidazole, pH 7.5). The suspension was incubated for 20 minutes, and the eluate was recovered. Two additional elutions were performed using 250 μL of a strong elution buffer (20 mM Tris-HCl, 0.5 M NaCl, 375 mM imidazole, pH 7.5), each followed by magnetic separation.
ii. Clarified *E. coli* lysate Suspension of magnetic nanoparticles was prepared by sonication of 2 mg of MNPs in 1.35 mL of a washing buffer (20 mM Tris-HCl, 0.5 M NaCl, 20 mM imidazole, pH 7.5) for 5 minutes in an ultrasonic bath. Subsequently, 250 μL of the clarified *E. coli* lysate containing the His6-tagged protein was added, resulting in a final volume of 1.6 mL. The mixture was incubated for 1 hour with gentle agitation. After incubation, the mixture was placed in a magnetic rack and allowed to sediment the MNPs. The supernatant was then removed. The first wash was performed by adding 200 μL of the washing buffer, mixing well, and incubating for 20 minutes. The magnetic rack was used again to remove the supernatant. This process was repeated for a second wash. For protein elution, three stages were performed. In the first elution, 200 μL of a weak elution buffer (20 mM Tris-HCl, 0.5 mol/L NaCl, 300 mmol/L imidazole, pH 7.5) was added, mixed well, and incubated for 20 minutes before removing the supernatant using the magnetic rack. In the second elution, a strong elution buffer (20 mM Tris-HCl, 0.5 M NaCl, 1 M imidazole, pH 7.5) was used following the same procedure. Finally, a third elution was performed with the same strong elution buffer.
iii. Large-scale clarified lysate For large-scale purification, 30 mg of MNPs were suspended in 13.5 mL of washing buffer (20 mM Tris-HCl, 0.5 M NaCl, 20 mM imidazole, pH 7.5) and sonicated for 5–10 minutes in an ultrasonic bath. Then, 4 mL of clarified lysate containing His6 protein were added, and the suspension was incubated for 1 hour under gentle agitation. Magnetic separation was performed using a custom tool consisting of a strong magnet affixed to the tip of a plastic rod. This magnetic bar was inserted into a clean glass culture tube, which was then immersed in a 50□mL *Falcon* tube containing the sample. In this configuration, the magnet did not come into direct contact with the solution, enabling efficient nanoparticle capture. After separation, the rod was removed, allowing resuspension of the nanoparticles in the next buffer without irreversible adhesion. The unbound fraction (~ □ 24mL) was recovered, and the MNPs were washed twice with 3□mL of washing buffer. Elution was performed in three sequential steps: 3□mL of weak elution buffer (300□mM imidazole), followed by two elutions with 3□mL of strong elution buffer (1□M imidazole), each with 20 minutes of incubation.

### Protein electrophoresis

To monitor the presence of the His6 protein during the purification process, sodium dodecyl sulfate polyacrylamide gel electrophoresis (SDS-PAGE) was performed. Briefly, 10□μL of each sample were mixed with 10 □ μL of 2X SDS-PAGE loading buffer (0.5 □ M Tris-HCl pH 6.8, 50% glycerol, 5% β-mercaptoethanol, 0.1% bromophenol blue, 2% SDS), and heated at 100°C for 5 minutes prior to loading.

Electrophoresis was carried out using discontinuous 10% SDS-PAGE gels. The stacking gel (0.125 M Tris-HCl pH 6.8) was prepared with 1.2□mL distilled water, 260□μL 30% acrylamide/bis solution, 500□μL 0.5□M Tris-HCl pH 6.8, 20□μL 10% SDS, 20 □ μL 10% ammonium persulfate (APS), and 2 □ μL TEMED. The resolving gel (0.375 M Tris-HCl pH 8.8) contained 1.9□mL distilled water, 1.7□mL 30% acrylamide/bis solution, 1.3 □ mL 1.5 □ M Tris-HCl pH 8.8, 50□μL 10% SDS, 50 □ μL 10% APS, and 5 □ μL TEMED.

Electrophoresis was performed using the Tris-Glycine running buffer. Gels were run at 120□V for 30 minutes (until the dye front reached the resolving gel), followed by 150 □ V for approximately 1.5 hours. Molecular weight markers used were *Marker* (*PB-L*), Spectra Multicolor Broad Range Protein Ladder (*Thermo Scientific*) and *PageRuler* (*Fermentas*).

After electrophoresis, gels were stained with Coomassie Brilliant Blue R-250 staining solution (2.5□g Coomassie in 450□mL methanol, 100□mL glacial acetic acid, and 450□mL distilled water), followed by destaining with a 45% methanol /10% acetic acid solution.

### Dialysis

Dialysis was performed using Molecular Porous membrane tubing with 3.5 and 10 kDa molecular weight cut-offs (*Spectra/Por*), placed in beakers containing 500 mL of buffer under constant agitation. To remove imidazole from the protein solution, dialysis was carried out against 10 mM Tris-HCl buffer (pH 7.4) for 18 hours, followed by two additional dialysis steps of 1 hour each.

## Results and discussion

### Selective Interaction of Nanoparticles with His-Tagged Proteins

To evaluate the binding affinity of nanoparticles to His6 proteins, we performed purification assays with prepurified proteins. Briefly, the chromatography purified protein was mixed with the MNPs; after incubation, two washing steps were performed, followed by three elutions using an imidazole-containing buffer. In all steps, a magnetic rack was used to separate the fraction bound to the MNPs from the supernatant. The obtained fractions were analyzed by loading them into a 10% polyacrylamide gel.

To evaluate the specificity of the magnetic nanoparticles for His-tagged proteins, bovine serum albumin (BSA), a ~66 kDa untagged protein, and a ~21 kDa histidine-tagged protein A (His6-SC-ProtA) were used. More than 90% of the BSA was recovered in the unbound fraction after incubation with magnetic nanoparticles (pass-through: PT), and no protein was observed in the eluate (Fig. 1A). In contrast, for His6-SC-ProtA, we observed that approximately 50% was recovered in the unbound fraction, while the remaining 50% was recovered in the eluates (Fig. 1B). This result confirms that the synthesized MNPs have selective affinity for histidine-tagged proteins, which stick to the MNPs and are then efficiently eluted with the buffers used.

**Fig 1.**
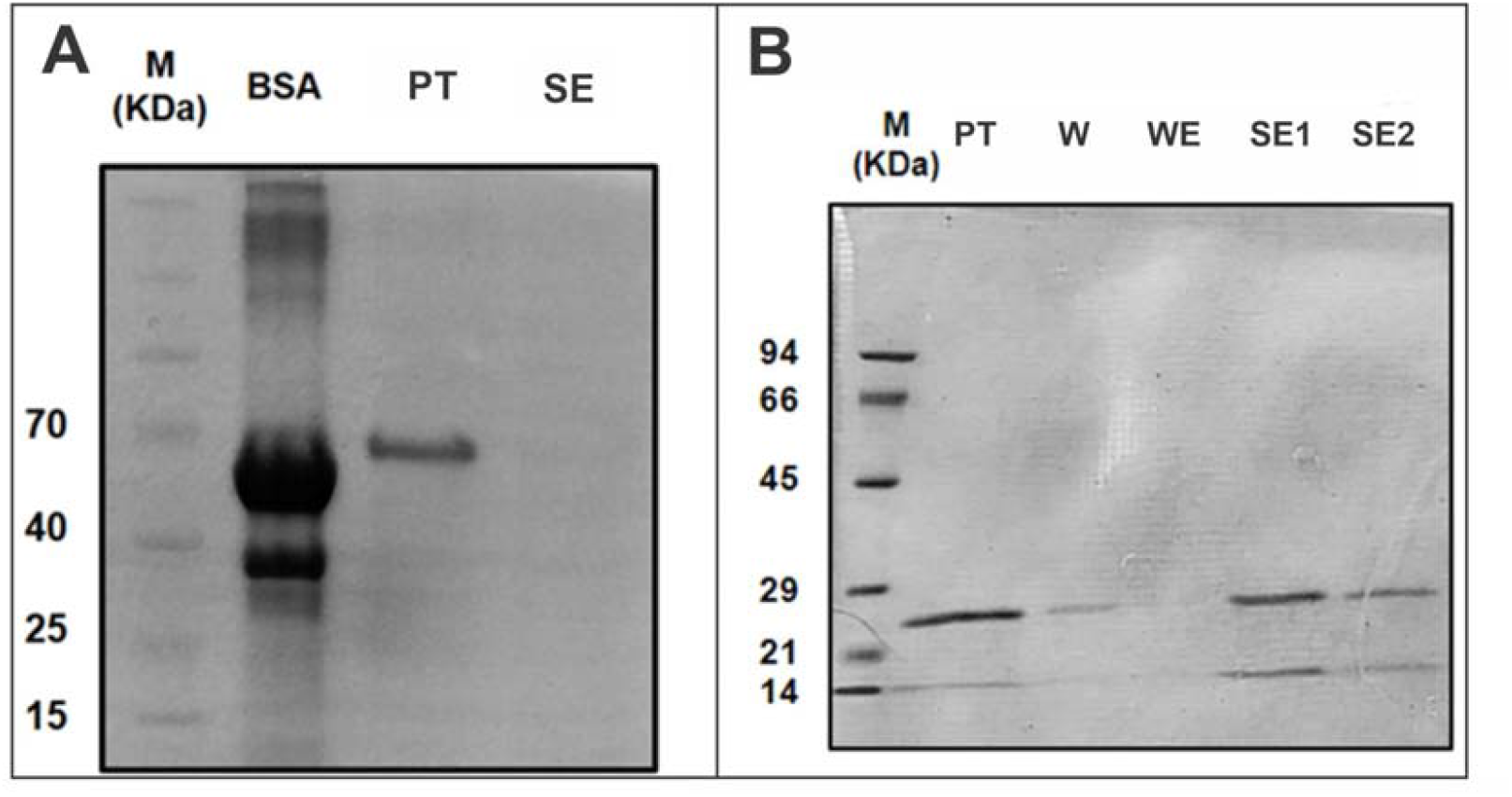
SDS-PAGE 10%. **A**. Bovine Serum Albumin (BSA) purification. **B**. His6-Sc-ProtA purification. M (kDa): protein *molecular weight markers (A: Spec*tra Multicolor Broad Range B: PB-L), PT: pass-through, W: wash, WE: weak elution, SE1: strong elution 1, SE2: strong elution 2.

### Isolation and Purification of Recombinant Proteins from *E. coli Lysate S*mall-Scale Purification

To test the performance of MNP@Ni^2+^ in the purification of His-tagged proteins from *E. coli* cultures, we induced the expression of His6-SC-ProtA and isolated it from 250 µL of lysate using magnetic separation. The purification was carried out according to the previously described protocol, employing two different elution buffers (300 and 1000 mM imidazole). SDS-PAGE was used to analyze all fractions (input, flow-through, washes, and elutions), enabling assessment of the recovery and purity of the His6 proteins (Fig. 2). We observed that His6-SC-ProtA (indicated by the arrow at ~21 kDa in the figure) was present in the bacterial lysate (L). Although a significant amount of the protein remained in the supernatant (PT) after magnetic affinity separation, the proportion of protein bound to the magnetic nanoparticles was sufficient to obtain purified protein in the elution fractions. Likewise, in the first wash (W1), we observed the presence of several proteins that appeared to be non-specifically bound to the MNPs, which were effectively removed in the second wash (W2), improving the purity of the eluted fractions. In the elution fractions a low-intensity band was observed corresponding to ProtA with 300 mM of imidazole (WE), this intensity increases considerably with the use of 1000 mM of imidazole (SE1 and SE2). It is important to highlight that a single, well-defined band is present in the elution fractions, confirming the specificity of the purification achieved through the selective interaction between the His-tag and the Ni^2+^-functionalized magnetic nanoparticles.

**Fig 2.**
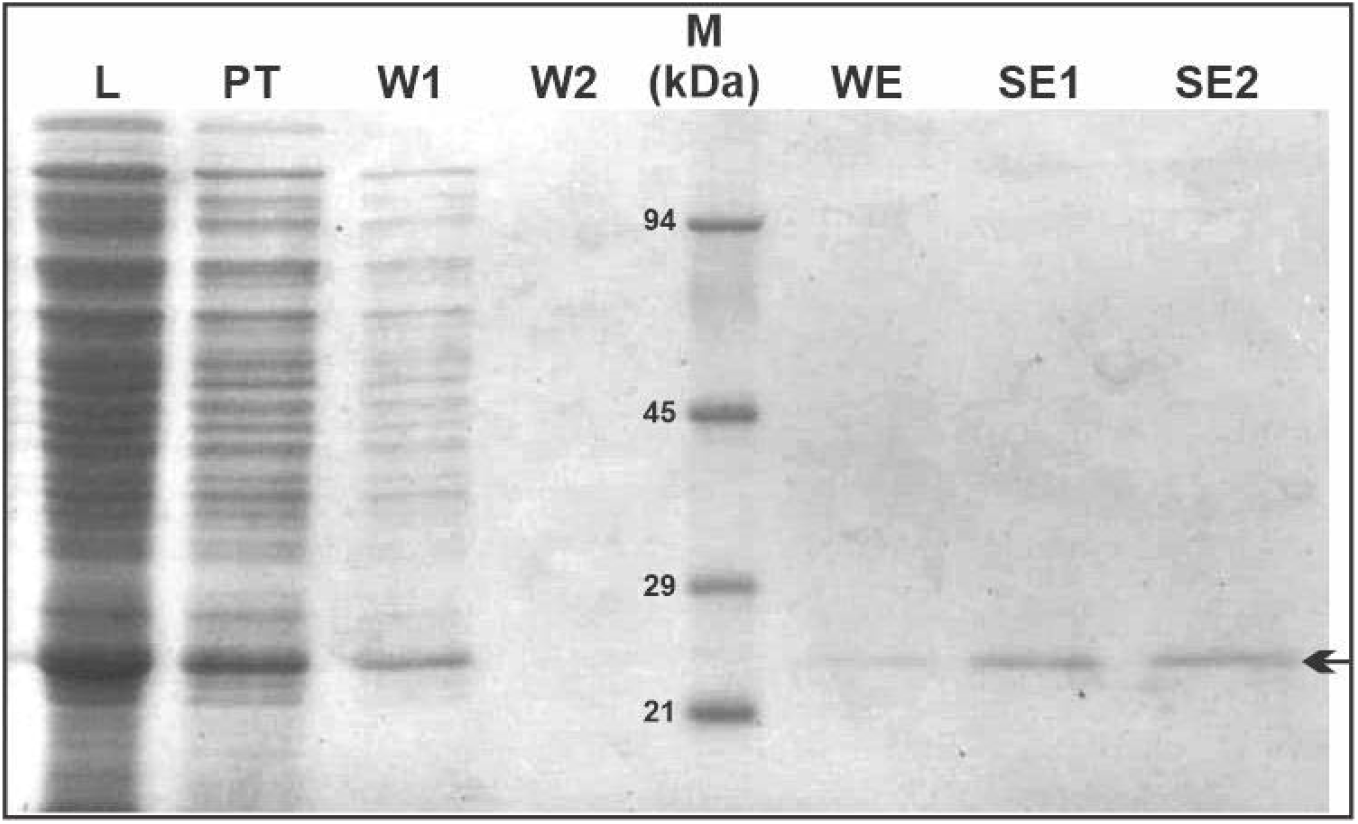
SDS-PAGE (10%) analysis of the purification steps of the His-tagged protein from *E. coli* lysate. Initial sample (L, clarified lysate), pass-through (PT), wash steps (W1 and W2), protein molecular weight marker (M), weak elution (WE), and strong elutions (SE1 and SE2). The arrow indicates the expected molecular weight of the target protein.

Using this protocol, we found that approximately 50% of the total His6-SC-ProtA present in the lysate binds to the MNP, while the remaining 50% is recovered in the pass-through (PT) fraction. Of the protein bound to the MNPs, about half is successfully recovered in the elution steps, with a small proportion lost during the wash steps. Importantly, the PT fraction can be subjected to a second round of incubation with MNPs to recover additional His6-SC-ProtA (Fig. S1). The protein concentration in the eluates was determined using *ImageJ* analysis, and the total amount of protein obtained (WE + SE1 + SE2) was expressed relative to the volume of lysate: 0.15 µg/µL of lysate. The yield obtained is comparable to those reported for standard affinity chromatography (Schmitt et al., 1993). Although direct comparisons with chromatographic methods are not strictly applicable due to the significantly smaller scale of our protocol, we highlight the efficiency of the magnetic nanoparticle-based purification system, which performs well even under low-volume conditions.

Similarly, we evaluated the capacity of the magnetic nanoparticles to purify other recombinant proteins of varying molecular weights, aiming to demonstrate the applicability of the system across a broad size range (approximately 20 to 100 kDa). Comparable results were obtained when purifying different recombinant proteins such as T7 RNA polymerase (~98 kDa), MMLV reverse transcriptase (~75 kDa), M protein (~60 kDa), and eIF4E protein (~45 kDa). In all cases, efficient purification was achieved, even when working with small-scale lysates (Fig. S2).

This section demonstrates the ability of MNP@Ni^2+^ to purify His-tagged recombinant proteins through magnetic separation and imidazole-mediated elution. The process proved to be specific and high-yielding, supporting the effectiveness of this method for efficient protein purification.

### Scaling Up: Yield Comparison with Chromatography Protein A

To obtain larger amounts of purified protein per assay, we designed a scaled-up purification protocol that is highly accessible and cost-effective (Fig. 3A). The volume of *E. coli* lysate used was 16 times greater than in the small-scale experiments (4 mL; see Materials and Methods section). We achieved excellent purification of the recombinant protein (Fig. 3B). Two washing steps were sufficient to remove non-specifically bound proteins, and the strong elution buffer (1 M imidazole) released the protein from the magnetic nanoparticles more efficiently than the mild elution buffer. With two strong elutions, it was possible to recover nearly all of the bound protein.

**Fig 3.**
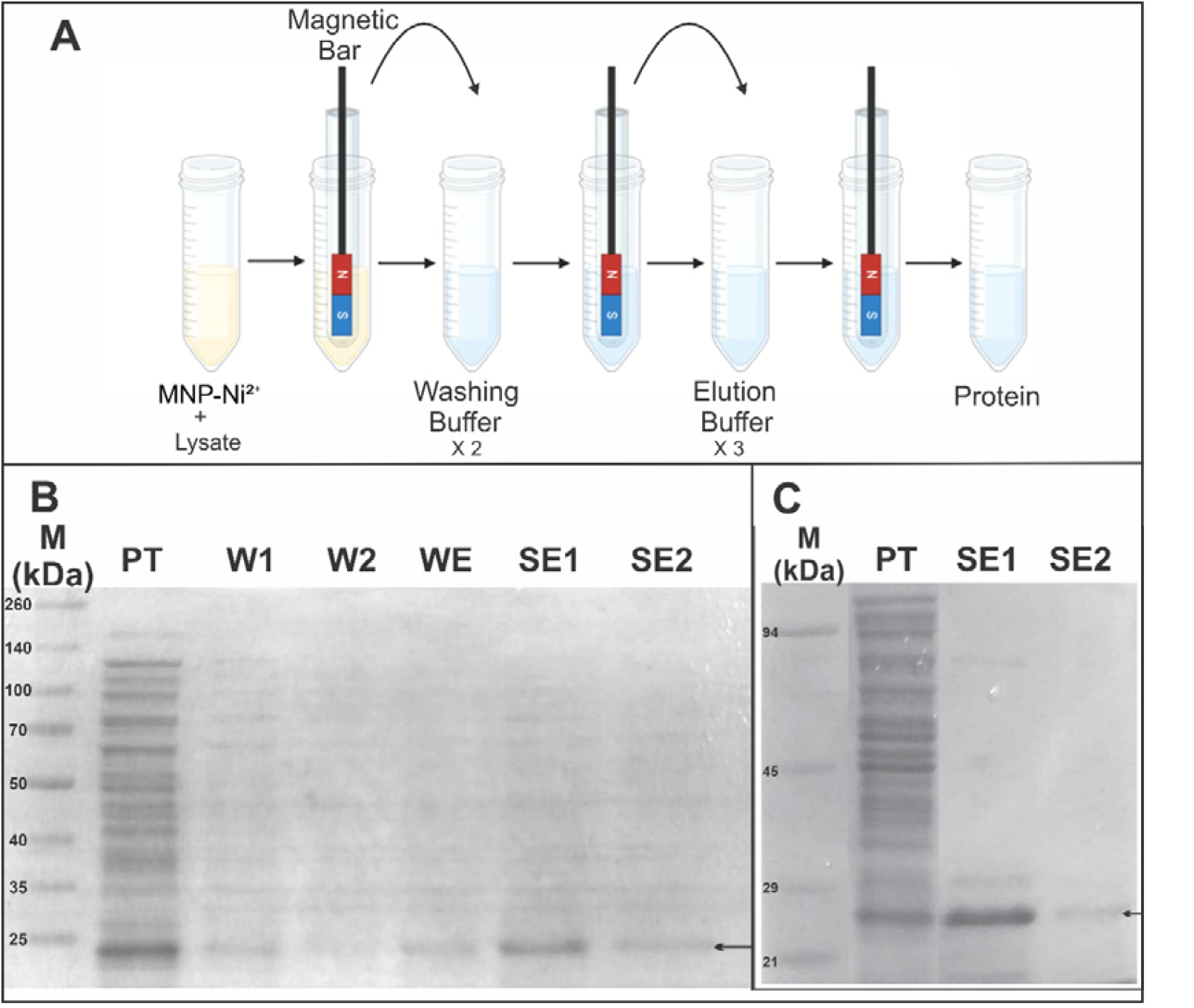
Protein purification using Ni^2+^ functionalized magnetic nanoparticles. **A**. Schematic representation of the purification workflow using a magnetic rod system. His6 proteins were bound to MNP@Ni^2+^ from clarified lysates, followed by two washes with buffer containing imidazole and three sequential elutions. Magnetic separation was performed using a bar magnet enclosed in a glass tube to enable efficient nanoparticle recovery and resuspension. **B**. SDS-PAGE analysis of fractions obtained during purification from E. coli lysate. **C**. SDS-PAGE of fractions from the same lysate purified using column chromatography (*AKTA Start*, 1⍰mL *HisTrap FF column, Cytiva*). Lane M: protein molecular weight marker; PT: pass-through; W1 and W2: first and second washes (20⍰mM imidazole); WE: weak elution (300⍰mM imidazole); SE1 and SE2: strong elutions (1 or 0.5⍰M imidazole). Arrows indicate the expected position of the target protein.

In order to compare the efficiency of our method with the gold standard, we purified the protein from the same lysate using column chromatography (*Akta Start-1ml His-Trap FF column, Cytiva*) (Fig. 3C). The magnetic nanoparticle-based method yielded a higher amount of purified protein compared to chromatography, a concentration of 0.3 mg of protein per mL of lysate was obtained using IMAC, while large-scale purification with MNPs yielded 0.6 mg/mL. This suggests approximately a two-fold increase in purification efficiency. However, it is important to note that the final elution volume was larger in the MNP-based method, which may require a subsequent concentration step. Nonetheless, such concentration steps—such as dialysis or centrifugal filters (e.g., Centricon)—are often also required following IMAC purification.

Some variation was observed in the amount of protein detected in the flow-through fractions, which may indicate saturation of the IMAC system. It is possible that using a column with greater binding capacity or volume could improve the overall yield and bring it closer to that obtained with the MNP-based method. Nevertheless, MNPs also exhibit saturation, as some protein was consistently detected in the nanoparticle-associated fractions that did not bind or elute efficiently. Despite this, the MNP-based approach proved to be an efficient, significantly simpler, and more cost-effective alternative to conventional chromatography, suitable for both analytical and preparative applications in the laboratory.

## Conclusions

MNPs capable of purifying His tagged recombinant proteins were designed and produced, demonstrating an efficiency comparable to the most commonly used purification methods. A magnetic nanoparticle-based platform was developed and validated for the efficient, selective, and scalable purification of His-tagged proteins. Ni^2+^-functionalized MNPs were synthesized and shown to exhibit high specificity for His6 proteins, as demonstrated by selective retention and imidazole-mediated elution from both pre-purified samples and complex *E. coli* lysates. The selective affinity for histidine residues was confirmed through comparative assays with bovine serum albumin and His6-SC-ProtA.

The method was successfully applied to purify various recombinant proteins ranging from 20 to 100 kDa, including T7 RNA polymerase, MMLV reverse transcriptase, M protein, and eIF4E. Small-scale purification protocols were optimized with minimal non-specific binding, allowing highly pure proteins to be obtained in a single elution step. When scaled up, yields approximately double those obtained with conventional immobilized metal affinity chromatography (IMAC) were achieved, while maintaining excellent purity.

The MNP-based purification protocol was found to be significantly more cost-effective and easy to implement, requiring no sophisticated equipment, thus making it highly suitable for laboratory-scale applications in both research and industrial settings. Further evaluation is required to assess the performance of these magnetic nanoparticles at preparative and productive scales for large-scale industrial purposes.

## Acknowledgment

This work was supported with grants from CONICET (PIP 11220210100751CO), AGENCIA I+D+I (PICT-2020-SERIEA-00865/PICT-2021-SERIEA-0034), and UNLP (11/X953).

## Supplemental Material

**Fig S1.**
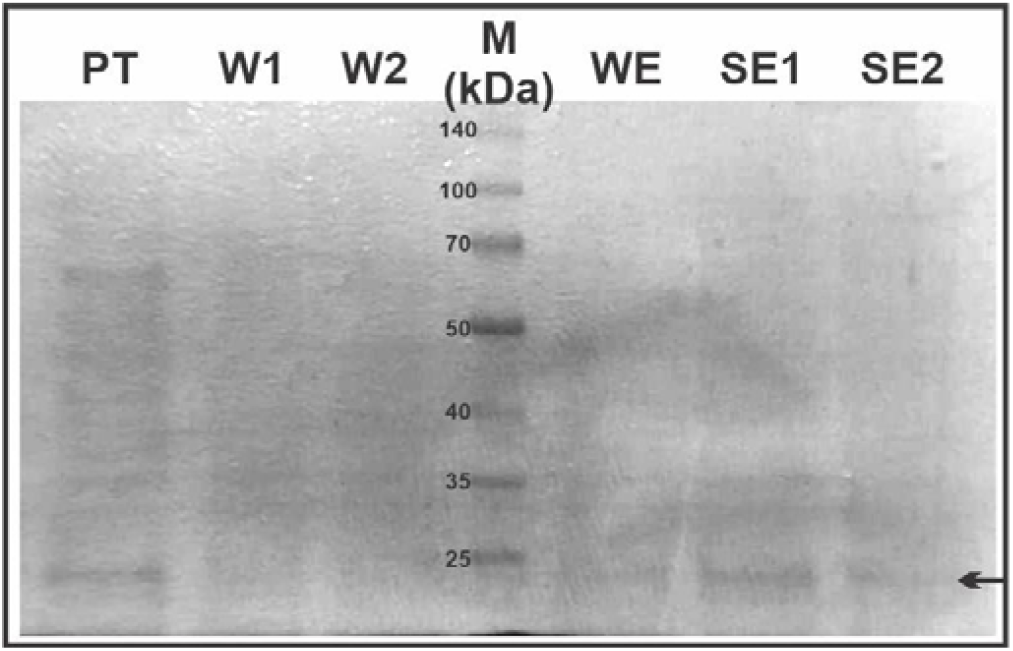
SDS-PAGE (10%) analysis of the second round of purification from pass-through. Initial sample: pass-through for His6-SC-PA purification with MNP@Ni^2+^ from *E.coli* lysate (PT), wash steps (W1 and W2), protein molecular weight marker (M), weak elution (WE), and strong elutions (SE1 and SE2). The arrow indicates the expected molecular weight of the target protein.

**Fig S2.**
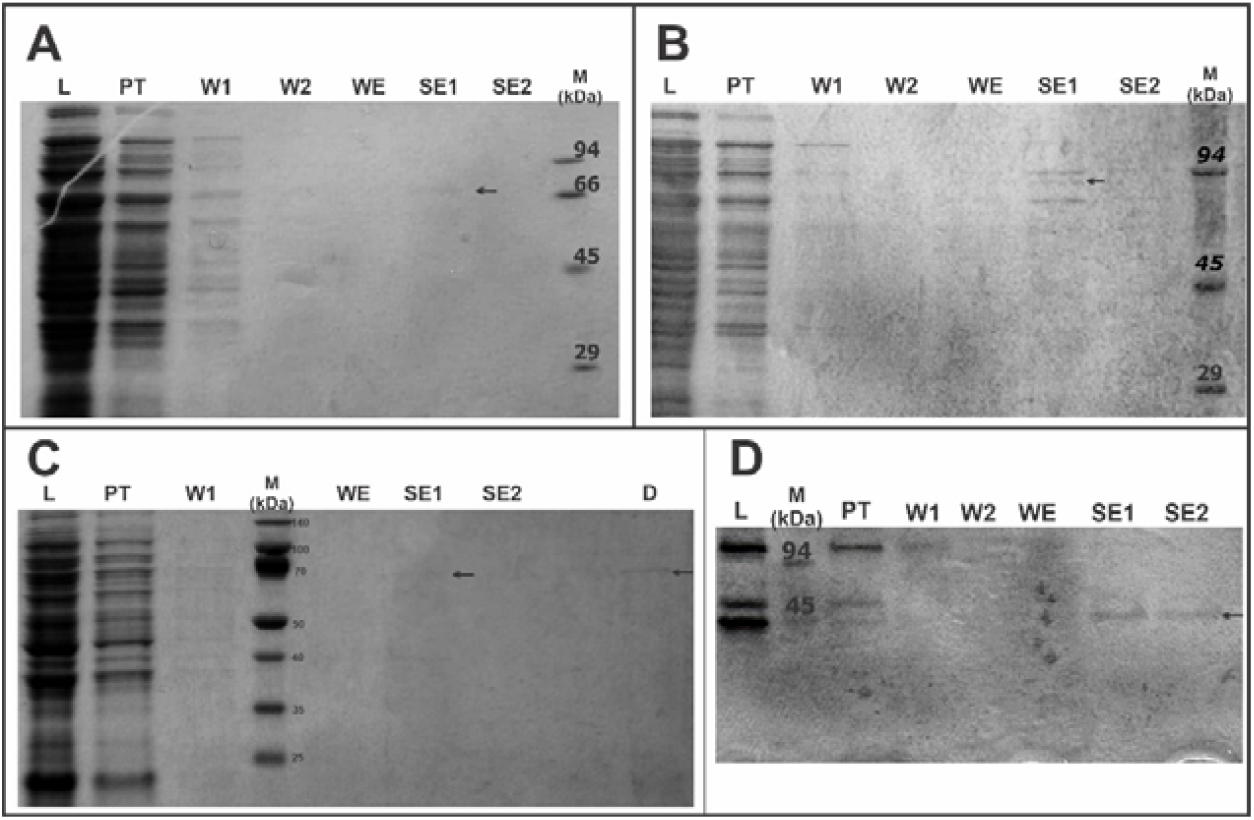
SDS-PAGE (10%) analysis of protein purification with MNP@Ni^2+^. **A**. M protein (~60 kDa). **B**. T7 RNA polymerase (~98 kDa). **C.** MMLV reverse transcriptase (~75 kDa). D. eIF4E protein (~ 45 kDa). Initial sample: *E.coli* lysate (L), pass-through (PT), wash steps (W1 and W2), protein molecular weight marker (M), weak elution (WE), strong elutions (SE1 and SE2) and dialyzed (D). The arrow indicates the expected molecular weight of the target protein.

