## Supplementary material for "Magnetic Affinity-Based Purification of His-Tagged Proteins Using Ni^2+^-Functionalized Nanoparticles": Fig S1 and Fig S2

### Supplemental Material

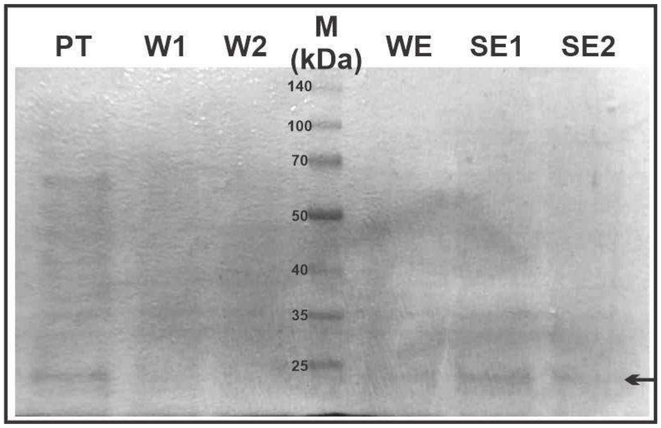


**Fig S1. SDS-PAGE (10%) analysis of the second round of purification from pass-through.** Initial sample: pass-through for His6-SC-PA purification with MNP@Ni$^{2+}$ from *E.coli* lysate (PT), wash steps (W1 and W2), protein molecular weight marker (M), weak elution (WE), and strong elutions (SE1 and SE2). The arrow indicates the expected molecular weight of the target protein.

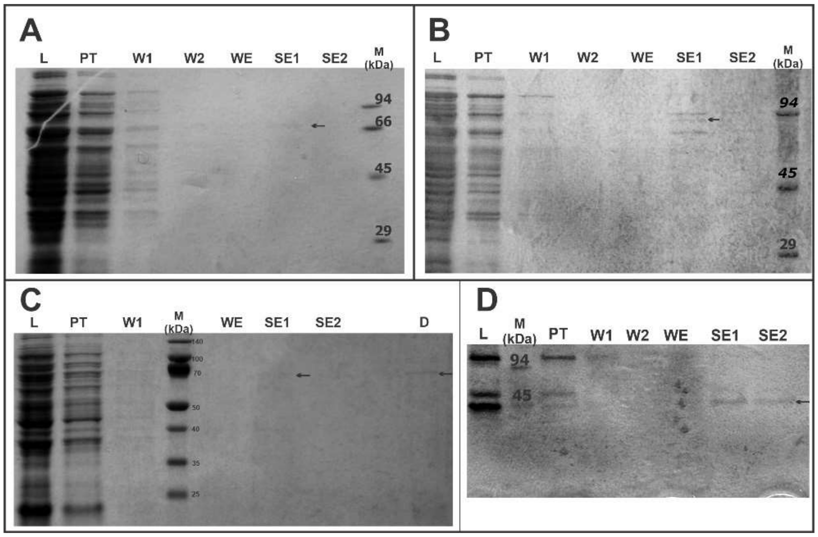


**Fig S2. SDS-PAGE (10%) analysis of protein purification with MNP@Ni$^{2+}$. A.** M protein (~60 kDa). **B**. T7 RNA polymerase (~98 kDa). **C**. MMLV reverse transcriptase (~75 kDa). D. eIF4E protein (~ 45 kDa). Initial sample: *E.coli* lysate (L), pass-through (PT), wash steps (W1 and W2), protein molecular weight marker (M), weak elution (WE), strong elutions (SE1 and SE2) and dialyzed (D). The arrow indicates the expected molecular weight of the target protein.
